# Glycogen-dependent demixing of frog egg cytoplasm at increased crowding

**DOI:** 10.1101/2021.04.11.439352

**Authors:** James F. Pelletier, Christine M. Field, Margaret Coughlin, Lillia Ryazanova, Matthew Sonnett, Martin Wühr, Timothy J. Mitchison

## Abstract

Crowding increases the tendency of macromolecules to aggregate and phase separate, and high crowding can induce glass-like states of cytoplasm. To explore the effect of crowding in a well-characterized model cytoplasm we developed methods to selectively concentrate components larger than 25 kDa from *Xenopus* egg extracts. When crowding was increased 1.4x, the egg cytoplasm demixed into two liquid phases of approximately equal volume. One of the phases was highly enriched in glycogen while the other had a higher protein concentration. Glycogen hydrolysis blocked or reversed demixing. Quantitative proteomics showed that the glycogen phase was enriched in proteins that bind glycogen, participate in carbohydrate metabolism, or are in complexes with especially high native molecular weight. The glycogen phase was depleted of ribosomes, ER and mitochondria. These results inform on the physical nature of a glycogen-rich cytoplasm and suggest a role of demixing in the localization of glycogen particles in tissue cells.

## Introduction

The concentration of macromolecules in cytoplasm is thought to reflect a balance between competing evolutionary pressures. Broadly speaking, increased concentrations tend to speed up biochemistry, but very high concentrations can cause excessive crowding, leading to deleterious interactions and potentially freezing of biochemistry that depends on dissociation reactions of macromolecules. In more precise terms, reaction-limited reaction rates increase with crowding due to increased re-collision frequency of reactants (Kim and Yethiraj, 2009). At the same time, diffusion-limited reaction rates increase with concentration but decrease with viscosity, and viscosity increases with concentration (Dill et al., 2011; van den Berg et al., 2017). Crowding tends to promote macromolecular assembly reactions, including linear polymerization and phase-separated condensate assembly (Andre and Spruijt, 2020). Increasing crowding may contribute to a liquid-to-glass transition in bacterial and yeast cytoplasm under energy stress (Joyner et al., 2016; Munder et al., 2016; Parry et al., 2014) or mechanical compression (Okumus et al., 2016). Whether this glass transition is a universal response to increased crowding is not known, though local dehydration was shown to decrease the mobility of macromolecules in human cells in tissue culture (Charras et al., 2009).

The concept of crowding is related to those of excluded volume effects (Rivas and Minton, 2016) and colloid osmotic pressure (Mitchison, 2019). Importantly, the ability of a molecule to exert crowding effects, and the response of molecules to crowding, depend on their size. Cytoplasm is polydisperse and soluble biomolecules cover orders of magnitude in radius, spanning from water molecules to ribosomes and other large complexes. Increased crowding tends to preferentially affect larger molecules because they start to physically interact before smaller molecules (Hwang et al., 2016). This reduces the mobility of large molecules, such as ribosomes, at concentrations which still allow smaller molecules to move in the crevices between them (Delarue et al., 2018). One might expect increased crowding to induce demixing of the largest components while leaving smaller components free to diffuse; however, the effect of crowding on cytoplasm is complicated by binding reactions, which may cause smaller molecules to demix along with larger ones to which they bind.

The storage polysaccharide glycogen is one of the most abundant macromolecules in many animal cells, on the order of 10% (w/w) in fed liver tissue (Dowler and Mottram, 1918; Prats et al., 2018); however, it is often forgotten in discussions of the structure and dynamics of cytoplasm. Glycogen is synthesized as a particle that consists of a highly branched, covalent polymer of glucose, usually initiated by polymerization from the surface of the nucleation protein glycogenin (Prats et al., 2018). Glycogen particles can be of different sizes, with a diameter of ∼20 nm typical (Ioan et al., 1999a, b). Glycogen particles are highly soluble in water, though they can form higher order assemblies (Nawaz et al., 2021; Prats et al., 2018). The cell biology and biophysics of glycogen have been little studied in recent years, though a recent report of the glycogen-binding proteome is relevant to this work (Stapleton et al., 2013; Stapleton et al., 2010).

*Xenopus* egg extracts provide a well-characterized model cytoplasm. They are prepared from eggs by centrifugal crushing with minimal dilution and maintain many of the biochemical and biophysical properties of native egg cytoplasm. Egg extracts contain ∼80 mg/mL protein and ∼80 mg/mL glycogen. The glycogen provides an energy store for the developing embryo and may also provide a crowding function that promotes assembly of nuclei (Hartl et al., 1994) and mitotic spindle poles (Groen et al., 2011). We set out to investigate the effect of changes in crowding on cytoskeleton dynamics in egg extract, but when we increased crowding by 1.4x or higher, we observed bulk demixing into two phases, which was unexpected. Here, we report that this demixing depends on glycogen and generates one phase that is highly glycogen-enriched. These observations probably do not model any normal egg biology, but they are informative concerning the physical properties of glycogen and its influence on those of the cytoplasm.

## Results

### Crowded *Xenopus* egg extracts exhibit liquid-liquid demixing

We developed two methods to selectively increase macromolecular crowding of *Xenopus* egg extracts while minimally perturbing ionic strength, pH and metabolite concentrations. Both methods gave similar results. In the first method, dry Sephadex G-25 gel filtration resin was added directly to extract and then removed a few minutes later by centrifugal filtration. As the resin swells, it absorbs water and small molecules, but macromolecules greater than 25 kDa are excluded and therefore become more crowded (Fig 1A). This method was convenient for small volumes of extract. The crowding factor depended on the amount of dry resin added per volume extract (Figs 1B, S1). In the second method, a 30 kDa MWCO centrifugal filter unit was used to concentrate macromolecules greater than 30 kDa. This method was convenient for larger volumes. The crowding factor was measured by adding a macromolecular fluorescent probe, such as Streptavidin (53 kDa) fused to Alexa Fluor 647 (Fig 1C), then comparing the fluorescence intensity of a specimen of fixed depth before and after crowding by fluorescence microscopy with a low magnification objective (Fig S1).

**Figure 1.**
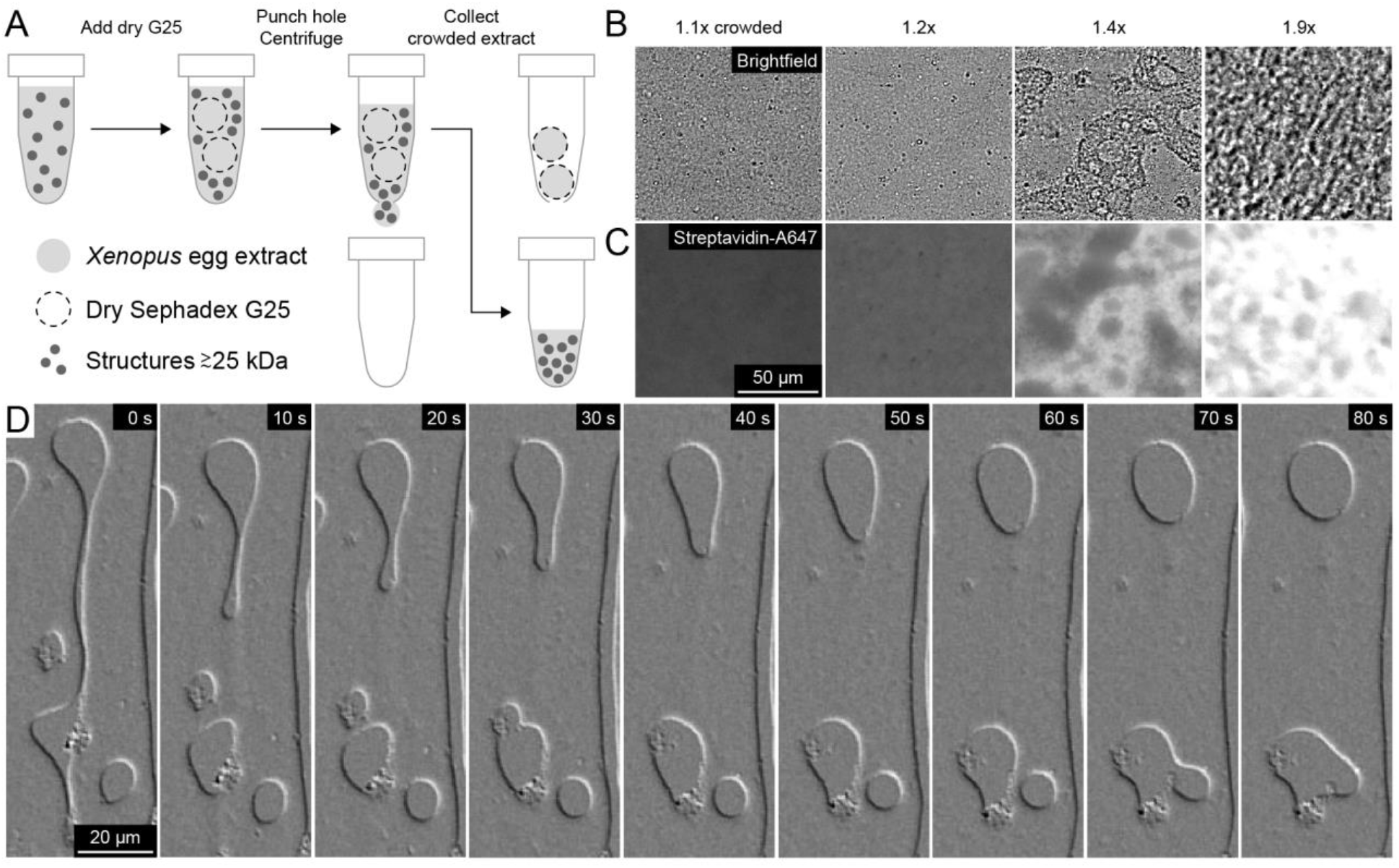
Crowded *Xenopus* egg extracts exhibit liquid-liquid demixing. (A) The macromolecular fraction greater than 25 kDa of *Xenopus* egg extracts was crowded by adding dry Sephadex G-25 resin, which selectively imbibes water and small molecules as it swells. (B) Brightfield images. Above 1.4x crowding, the extract remained liquid-like and exhibited patches with different light scattering properties. Above 1.9x crowding, precipitation occurred. (C) The crowding factor was estimated by quantification of fluorescence of a protein probe greater than 25 kDa added before crowding, in this case Streptavidin (53 kDa) fused to Alexa Fluor 647. The Streptavidin-A647 partitioned into one of the phases. Crowding factors estimated from lower magnification images in Fig S1. (D) Time lapse differential interference contrast (DIC) images of a 1.4x crowded extract confined between coverslips. The phases exhibited hallmarks of liquids, including deformation under flow, splitting, fusion, and rounding by surface tension. See Video S1.

When crowding was increased to 1.2x, the extract remained similar in appearance to uncrowded extracts, as observed in brightfield and fluorescence images (Fig 1B,C). At 1.4x-1.5x, the extract underwent spontaneous demixing over a few minutes at both 0 °C and 20 °C. This unexpected phenomenon was further characterized. At 1.9x the extract appeared to precipitate, and this regime was not examined further. The phases formed throughout the sample and remained co-mingled in the tube. They had different densities and could be separated in bulk by centrifugation at 20000 rcf for 20 min. The volumes of the two phases after centrifugation were similar. The denser phase had a higher index of refraction of 1.39, and the less dense phase had a lower index of refraction of 1.38. This refractive index difference made demixing easy to follow by phase contrast or DIC microscopy. Both phases exhibited all the hallmarks of liquids, including deformation under flow, splitting, fusion, and rounding towards a spherical shape driven by surface tension (Hyman et al., 2014) (Fig 1D, Video S1).

### Role of glycogen in demixing

The higher refractive index of the denser phase suggested non-equal distribution of glycogen, which has a higher density and refractive index than protein. We measured the glycogen concentration in unperturbed extract and the two phases using an assay that digested it to glucose for colorimetric quantification. Glycogen was highly enriched in the denser phase (Fig 2A). The glycogen concentration was 80 mg/mL in uncrowded crude extracts, 20 mg/mL in the less dense phase, and 250 mg/mL in the denser phase (Fig 2A) (Methods). Total protein was slightly enriched in the less dense phase (Fig 2B).

**Figure 2.**
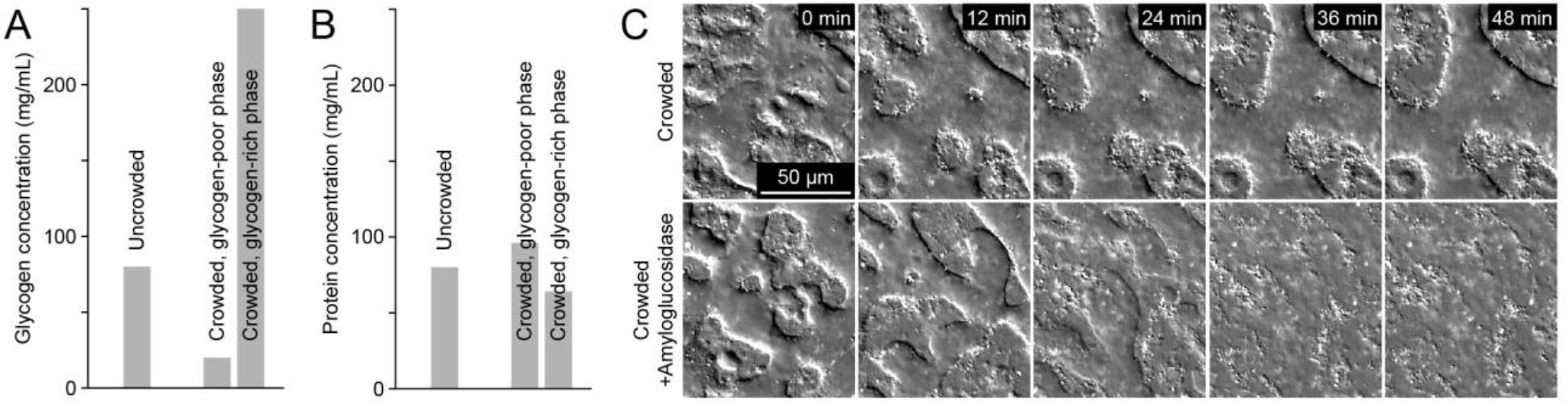
Role of glycogen in demixing. (A) Glycogen highly partitioned between the phases, as measured by a colorimetric assay. The concentration of glycogen in uncrowded extract was 80 mg/mL. In crowded extracts, the glycogen concentration was 20 mg/mL in the less dense phase and 250 mg/mL in the denser phase. (B) The less dense, glycogen-depleted phase had a higher protein concentration than the denser, glycogen-enriched phase. (C) DIC images. Top row: Control crowded extract remained demixed. Bottom row: Addition of amyloglucosidase (AG) after demixing caused the phases to dissolve and the system to return to a single phase.

To test for a role of glycogen in demixing, we hydrolyzed it using the enzyme amyloglucosidase (AG) (Methods). Glycogen digestion by AG blocked demixing when added before crowding (not shown) and reversed it when added after crowding, so demixing depended on glycogen (Fig 2C).

### Ultrastructure of glycogen-enriched (G) and -depleted (R) phases

Thin-section electron microscopy with conventional heavy metal staining was used to probe the ultrastructure of the phases. Uncrowded crude extracts appeared mottled with uniformly distributed mitochondria and ER (Fig 3A). Crowded extracts exhibited at least two major phases as in optical micrographs (Fig 3B). One of the phases had higher electron density than the other (Fig 3B). To identify the phases in electron micrographs, the phases were isolated in bulk by centrifugation then imaged separately (Methods). The glycogen-depleted phase (Fig 3C) had higher electron density than the glycogen-enriched phase (Fig 3D). Mitochondria concentrated at the interface between the phases (Fig 3B,E) and were also present within the glycogen-depleted phase (Fig 3C). The higher electron density, glycogen-depleted phase was textured with structures ∼25 nm in diameter, which we interpret as ribosomes (Fig 3E’’). Hereafter, we refer to the glycogen-depleted phase as “R” for ribosomes and the glycogen-enriched phase as “G” for glycogen.

**Figure 3.**
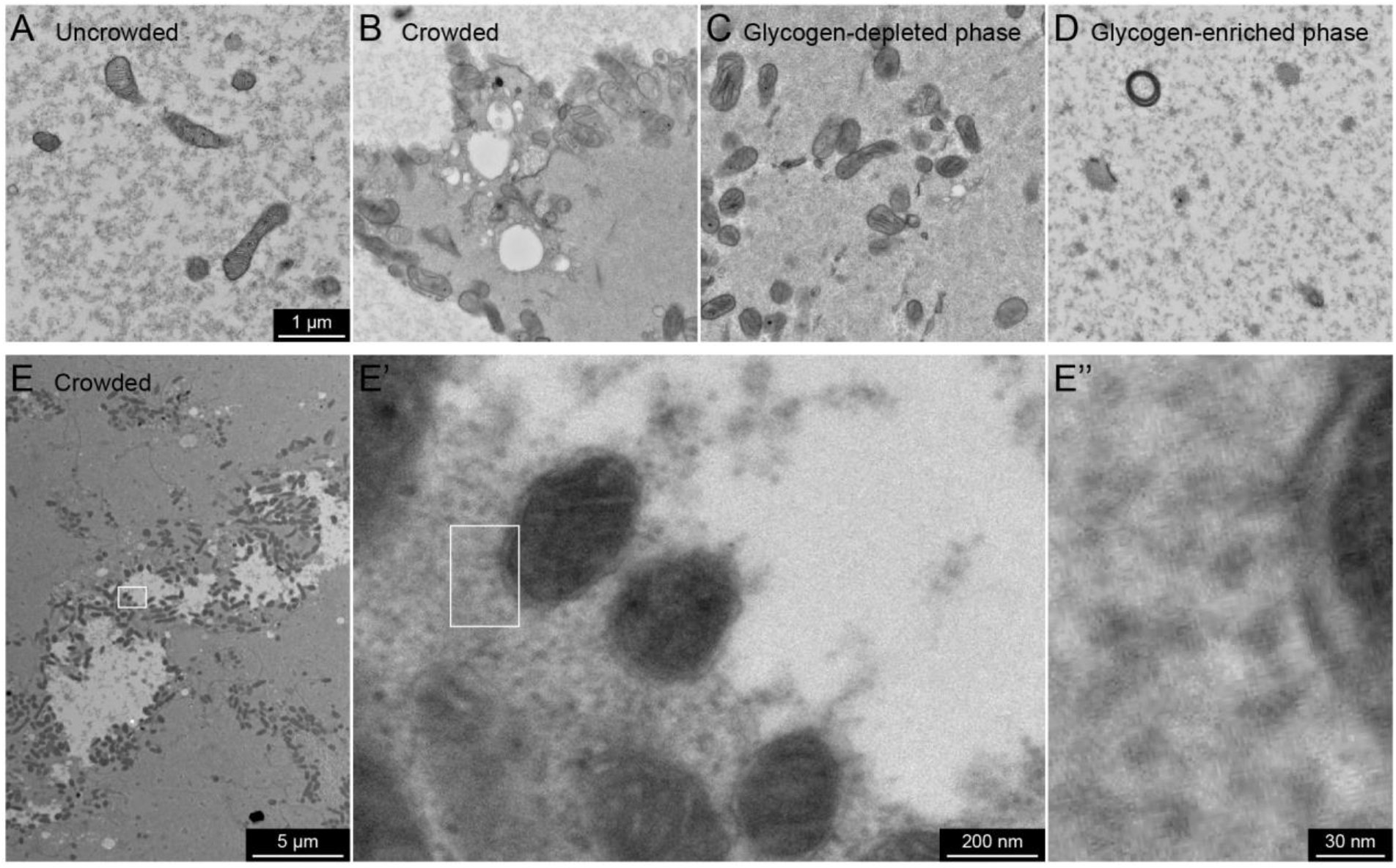
Ultrastructure of glycogen-enriched (G) and -depleted (R) phases. (A) Uncrowded crude extract did not exhibit bulk demixing but exhibited a mottled appearance. (B) Crowded extracts exhibited bulk demixing, with one phase higher electron density than the other. (C,D) The glycogen-depleted and glycogen-enriched phases were isolated from one another in bulk by centrifugation and imaged separately. The glycogen-depleted phase was higher electron density than the glycogen-enriched phase. (E) Mitochondria often localized along the interface between the phases or in the higher contrast phase. E’ is a zoom of the box in panel E, and E’’ is a zoom of the box in panel E’. The higher contrast phase had a granular appearance with features ∼25 nm in diameter, which we interpret as ribosomes.

### Fluorescent probes partitioned between G and R phases

Fluorescence microscopy provided a convenient method to observe the two phases and estimate the partition coefficient of macromolecules. Fluorescent probes such as EB1-mApple (57 kDa) and Streptavidin (53 kDa) labeled with Alexa Fluor 647 were added to crowded extracts. Then, to estimate partitioning of each probe between the phases, the G and R phases were isolated from one another in bulk by centrifugation (Methods). We could thus estimate the partition coefficients of the fluorescent probes (Fig 4A,B), as well as identify the mixed phases using the fluorescent probes (Fig 4C). Partition coefficients in mixed phases were similar to those in bulk (Fig 4D). Most probes partitioned preferentially into the R phase, including EB1-GFP (57 kDa), Fab fragment antibody (50 kDa) labeled with Alexa Fluor 647, and 70 kDa dextran-Alexa Fluor 488 (Fig 4E-G, I-J). Glycogen phosphorylase A (PYGL, 188 kDa as dimer), a glycogen-binding protein, labeled with Pacific Blue partitioned preferentially into the G phase (Fig 4K). Mitochondria imaged by NADH autofluorescence localized along the interface between phases (Fig 4H), as seen by electron micrographs (Fig 3B,E).

**Figure 4.**
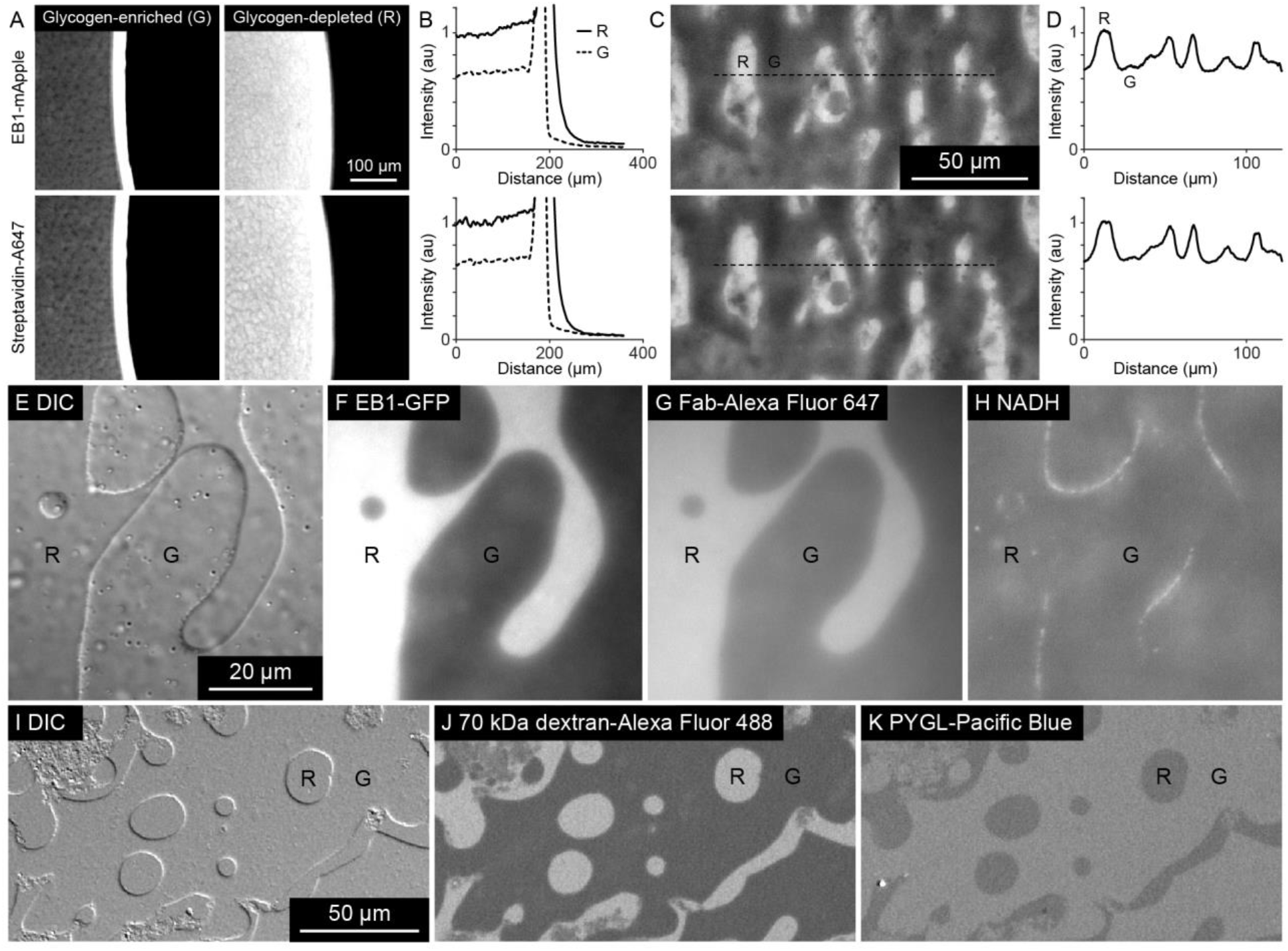
Fluorescent probes partitioned between G and R phases. (A) EB1-mApple and Streptavidin-A647 in G and R phases, isolated from one another in bulk by centrifugation. (B) Both probes partitioned preferentially into the R phase than the G phase. (C) Mixed phases in crowded extract. (D) Intensity profile along the black dotted lines overlaid on panel C. (E-H) EB1-GFP and Fab-Alexa Fluor 647 likewise partitioned preferentially into the R phase, while mitochondria imaged by NADH autofluorescence localized along the interface between phases, as seen by electron micrographs. (I-K) 70 kDa dextran-Alexa Fluor 488 partitioned into the R phase, while PYGL-Pacific Blue partitioned into the G phase.

### Protein partitioning depends on glycogen binding and native molecular weight (MW)

To quantify the proteomes of the G and R phases, we performed multiplexed mass spectrometry analysis using the MultiNotch MS3 method (Gupta et al., 2018; McAlister et al., 2014; Sonnett et al., 2018). For this analysis, we compared two methods for crowding the extract, using either Sephadex G-25 resin or 30 kDa MWCO centrifugal filter units. Results were similar for the two methods (Fig S2, Table S1). Fig 5 reports measurements averaged across the two methods and several repeats.

**Figure 5.**
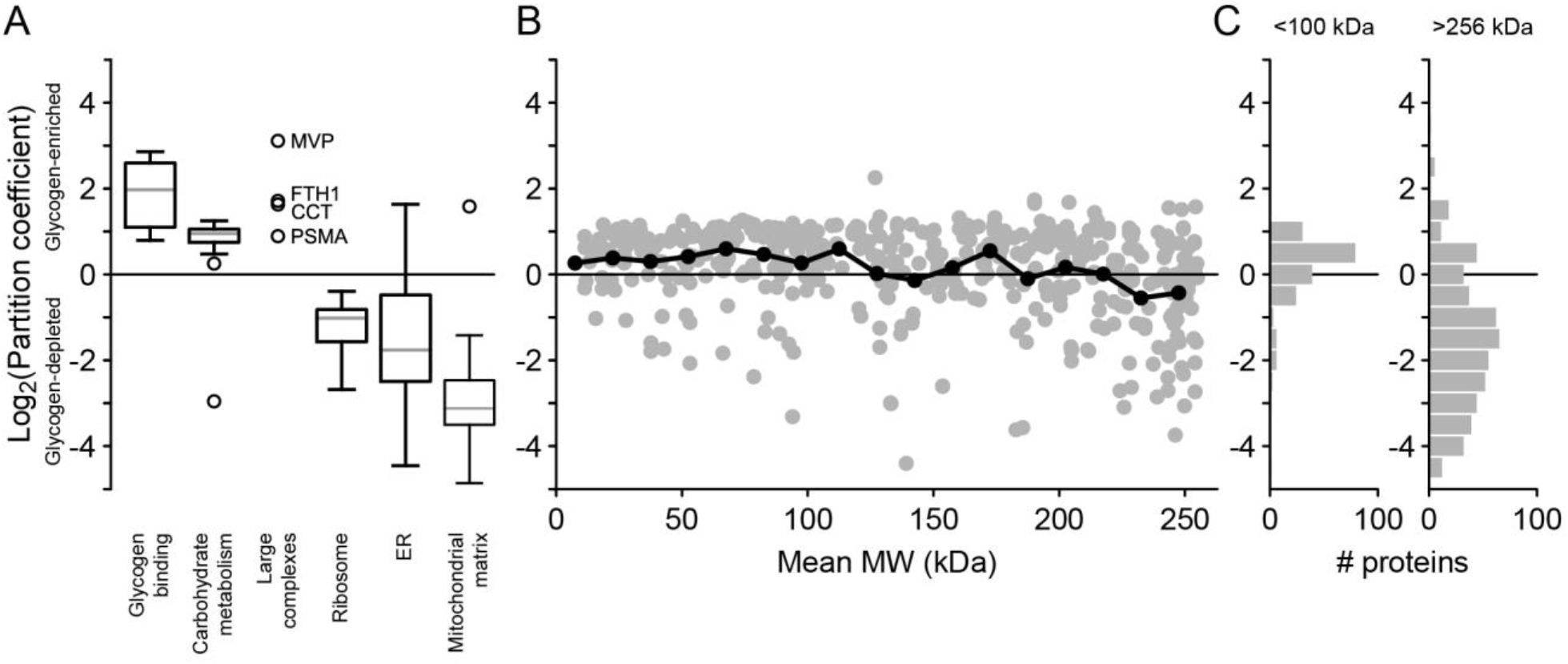
Proteomics analysis suggests partitioning depends on binding and native MW. (A) Log base 2 partition coefficients of proteins sorted by Gene Ontology terms. Gray lines represent median values, boxes represent first and third quartile values, whiskers show the range of the data up to 1.5x the interquartile range, and circles represent outliers. (B) Log base 2 partition coefficients with respect to native MW less than 256 kDa. (C) Log base 2 partition coefficients of all proteins in complexes with native MW less than 100 kDa or greater than 256 kDa. Replicates in Figure S2.

Known glycogen-binding proteins partitioned into the G phase, with log base 2 partition coefficients of 1-3 (Fig 5A). These values approach the partition coefficient of glycogen itself (Fig 2A). Ribosomal subunits, ER and mitochondrial proteins selectively partitioned into the R phase (Fig 5A), consistent with the electron micrographs (Fig 3C,E). Enzymes involved in carbohydrate metabolism also partitioned into the G phase, though with lower partition coefficients than proteins known to bind glycogen (Fig 5A). Several of the highest native MW protein complexes other than ribosomes partitioned into the G phase, such as major vault protein (MVP, 13 MDa), ferritin (FTH1, 450 kDa), chaperonin-containing T-complex (CCT, 960 kDa), and the 26S proteasome (PSMA, 2 MDa) (Fig 5A). We then plotted log base 2 partition coefficients with respect to native MW (Fig 5B), based on previous estimates of native MW with an upper bound of 256 kDa (Wühr et al., 2015). Proteins with native MW smaller than 100 kDa had a slight preference for the G phase, with average log base 2 partition coefficient 0.4 ± 0.8 (Fig 5C). In contrast, most proteins with native MW greater than 256 kDa partitioned into the R phase, with average log base 2 partition coefficient -1.3 ± 1.6 (Fig 5C).

## Discussion

When macromolecular crowding of *Xenopus* egg extracts was increased 1.4x over control, we observed glycogen-dependent demixing into liquid G and R phases that were enriched in glycogen and ribosomes, respectively. This was unexpected, since reports from other systems led us to expect a transition to a glass-like state. We suspect liquid-liquid demixing is promoted by the high glycogen concentration in egg cytoplasm, which is comparable to that in hepatocytes from fed liver.

Mass spec analysis suggested both binding and native MW contributed to partitioning of proteins between the phases. In terms of binding, glycogen-binding proteins partitioned preferentially into the G phase with an average partition coefficient similar to that of glycogen itself (Fig 5A). Enzymes involved in carbohydrate metabolism also partitioned into the G phase (Fig 5A). These spanned a range of native MWs from 33 to 241 kDa, with no apparent correlation between partition coefficient and native MW (Table S1). Some of these have been shown to bind glycogen in an adipocyte glycogen proteome (Stapleton et al., 2013; Stapleton et al., 2010). Association between glycolytic enzymes has also been reported, but its functional significance remains unclear (Schmitt and An, 2017). Consistent with a role for native MW, the G phase enriched several especially large protein complexes not known to bind glycogen (Stapleton et al., 2013; Stapleton et al., 2010) (Fig 5A). Relative contributions of binding and entropic considerations may be considered within excluded volume theory (Rivas and Minton, 2016).

Frog eggs evolved in a freshwater environment and *Xenopus* eggs are not known to exhibit desiccation resistance. Thus, the demixing we observed is unlikely to be of direct physiological relevance; however, it may provide clues to glycogen organization in tissue cells. Glycogen particles often appear as aggregates in multiple animal tissues (Coimbra and Leblond, 1966; Galavazi, 1971; Porter and Bruni, 1959; Revel, 1964; Revel et al., 1960; Sheldon et al., 1962) and chloroplasts (Crumpton-Taylor et al., 2012; Kasperbauer and Hamilton, 1984). Fawcett (1981) summarized extensive EM studies as showing that “glycogen is seldom uniformly distributed in the cytoplasm but tends to accumulate in dense regional deposits.” Our results suggest that physical demixing may contribute to high local concentrations of glycogen, though we cannot rule out other mechanisms including binding interactions between glycogen particles and local concentration of biosynthetic enzymes.

Our observations are also relevant to *Xenopus* egg extract technical considerations. Concentration of extract using 100 kDa filtration units was shown to increase the stability of extracts to freeze-thaw cycles (Takagi and Shimamoto, 2017). Those authors used ∼1.2x crowding, which is just below the concentration factor needed for demixing. Exploration in the crowded but still mixed regime may facilitate study of the effect of crowding on biochemical processes. Recent work examined how crowding affects microtubule polymerization using osmotic perturbation of fission yeast (Molines et al., 2020). It will be interesting to ask similar questions in cytoplasmic extracts.

## Materials and Methods

### Preparation of Xenopus egg extracts

*Xenopus* egg extracts were prepared as described previously (Field et al., 2017). Most experiments used extracts prepared with Cytochalasin D to prevent F-actin polymerization, in which case 100 µg/mL Cytochalasin D was added before the crushing spin at 18 °C, and 10 µg/mL Cytochalasin D was added after the crushing spin. Extracts with intact F-actin also demixed at similar crowding factors.

### Crowding of Xenopus egg extracts

*Xenopus* egg extracts were crowded by two methods, using Sephadex G-25 resin or 30 kDa MWCO filter units, which gave similar results. To crowd using coarse Sephadex G-25 gel filtration resin (Sigma-Aldrich Cat#GE17-0034-01), 30 µg dry resin was added to 150 µL extracts in a PCR tube. The resin was submerged and dispersed with a pipette tip, then the slurry was incubated for 5 min on ice. Then several holes were punched in the bottom of the PCR tube using the tip of a 27G needle, which makes holes small enough to retain resin in the tube. Then the PCR tube was placed inside a 0.5 mL tube, which was in turn placed inside a 1.5 mL tube for centrifugation. The tubes were centrifuged at 4000 rcf for 4 min to collect the crowded extract in the 0.5 mL tubes. To crowd using filter units, extracts were centrifuged in Amicon filter units with 30 kDa MWCO (Millipore #UFC5030BK). Crowded extracts were then stored on ice. To estimate crowding factors, macromolecular fluorescent probes were added to extracts before crowding.

### Fluorescent probes

To observe the two phases and estimate partitioning of fluorescent probes, we imaged Fab fragment antibody labeled with Alexa Fluor 647 (Jackson ImmunoResearch #111-607-003), Streptavidin labeled with Alexa Fluor 647 (Jackson ImmunoResearch #016-600-084), and Phosphorylase A (Sigma-Aldrich #P1261) labeled with Pacific Blue (Thermo Fisher #P10163).

### Digestion of glycogen

Amyloglucosidase (AG) (Sigma-Aldrich #A7420) was added to extracts to a final concentration of 1.25 mg/mL (8.7 µM).

### Measurement of glycogen concentration

The glycogen concentration was measured by a colorimetric assay based on the peroxidase sensitive dye 3,3′,5,5′-tetramethylbenzidine (TMB) (Sigma-Aldrich #860336). Glycogen was digested to glucose using amyloglucosidase (AG), then hydrogen peroxide was generated from glucose using glucose oxidase (Sigma-Aldrich #G7016), then TMB was oxidized to TMB diimine by the hydrogen peroxide with horseradish peroxidase (Sigma-Aldrich #P6782). In particular, reactions included 100 µg/mL TMB, 250 µg/mL AG, 250 µg/mL GO, 125 µg/mL HRP in 100 mM sodium citrate pH 5.0. Extracts and glycogen standards were titrated into reactions, and the TMB diimine absorbance at 660 nm was measured on a Synergy H1 plate reader (BioTek).

### Measurement of protein concentration

Protein concentration in each phase was measured by Micro BCA following TCA precipitation.

### Isolation of phases by centrifugation

Crowded extracts were centrifuged at 20000 rcf for 20 min. After centrifugation, the less dense R phase was aspirated into an 18 G blunt needle, carefully as not to disturb the interface between the R and G phases. The R phase appeared as two opaque layers of slightly different colors and both these were included in the R sample. Then a hole was punched in the bottom of the tube, and by pressing the top of the tube, the higher density G phase was pushed through the hole, likewise carefully as to avoid the interface between the phases.

### Electron microscopy

Extract was spread on coverslips then samples were prepared by standard methods. Extract samples were fixed with 1.5% glutaraldehyde in 0.1 M cacodylate buffer pH 7.4, post fixed with 1% osmium tetroxide/potassium ferrocyanide, en block stained with 1% uranyl acetate, dehydrated and embedded in Epon Araldite, then sectioned and on grid stained with uranyl acetate and lead citrate. Samples were viewed on a Tecnai G-2 BioTwin electron microscope and imaged with an AMT CCD camera.

### Mass spectrometry

Samples were denatured in 5 M guanidine thiocyanate, 5 mM dithiothreitol (DTT) (US Biological #D8070) for 10 min at 60 °C, then cysteines were alkylated with N-ethylmaleimide (NEM). The eluate was precipitated with trichloroacetic acid then subjected to proteolysis followed by the MultiNotch MS3 method as described (Gupta et al., 2018; McAlister et al., 2014; Sonnett et al., 2018), with channels normalized by the total number of counts. Native MWs were based on previous estimates (Wühr et al., 2015). Gene Ontology terms used for Fig 5A were Glycogen binding: GO:0005978 (Glycogen biosynthetic process, BP) and GO:0005980 (Glycogen catabolic process, BP); Carbohydrate metabolism: GO:0005975 (Carbohydrate metabolic process, BP) excluding GO:0005978 (Glycogen biosynthetic process, BP), GO:0005980 (Glycogen catabolic process, BP), GO:0005739 (Mitochondrion, CC), and GO:0005759 (Mitochondrial matrix, CC); Ribosome: GO:0005840 (Ribosome, CC); ER: GO:0005783 (Endoplasmic reticulum, CC); and Mitochondria matrix: GO:0005759 (Mitochondrial matrix, CC) excluding GO:0005829 (Cytosol, CC).

## Acknowledgements

This work was supported by NIH grants R35GM131753 (TJM) and R35GM128813 (MW), and MBL fellowships from the Evans Foundation, MBL Associates, and the Colwin Fund (TJM and CMF). JFP was supported by the Fannie and John Hertz Foundation, the Fakhri lab at MIT, the MIT Department of Physics, and the MIT Center for Bits and Atoms. The authors thank Keisuke Ishihara for critical feedback on the manuscript, the Nikon Imaging Center at Harvard Medical School and Nikon at MBL for imaging support, and the National Xenopus Resource at MBL for support. The EB1-GFP construct was a gift from Kevin Slep (UNC Chapel Hill, NC).

## Supplementary figures

**Figure S1.**
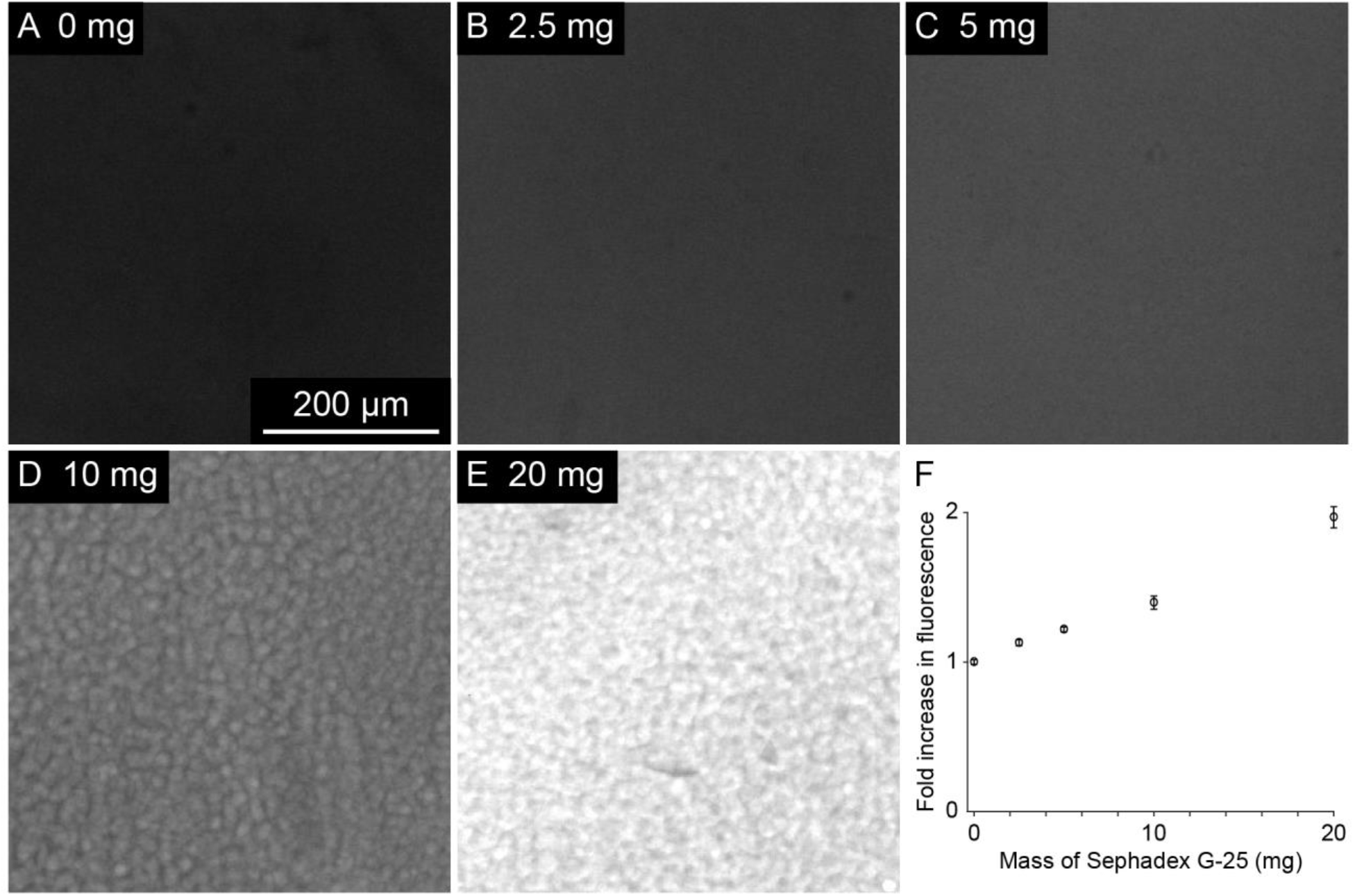
Estimation of crowding factors by measuring fold increase in fluorescence of a macromolecular probe. (Related to Fig 1B) (A-E) Streptavidin-Alexa Fluor 647 was added to extract, then the extract was split into 60 µL volumes. To vary the crowding factor, different amounts of dry Sephadex G-25 resin were added to each volume. After crowded extracts were collected (Methods), samples of fixed depth were prepared and imaged by fluorescence microscopy with a low magnification objective. (F) Fold increase in fluorescence with respect to mass of dry Sephadex G-25 resin per 60 µL extract. Error bars represent standard deviation of pixel intensity values.

**Figure S2.**
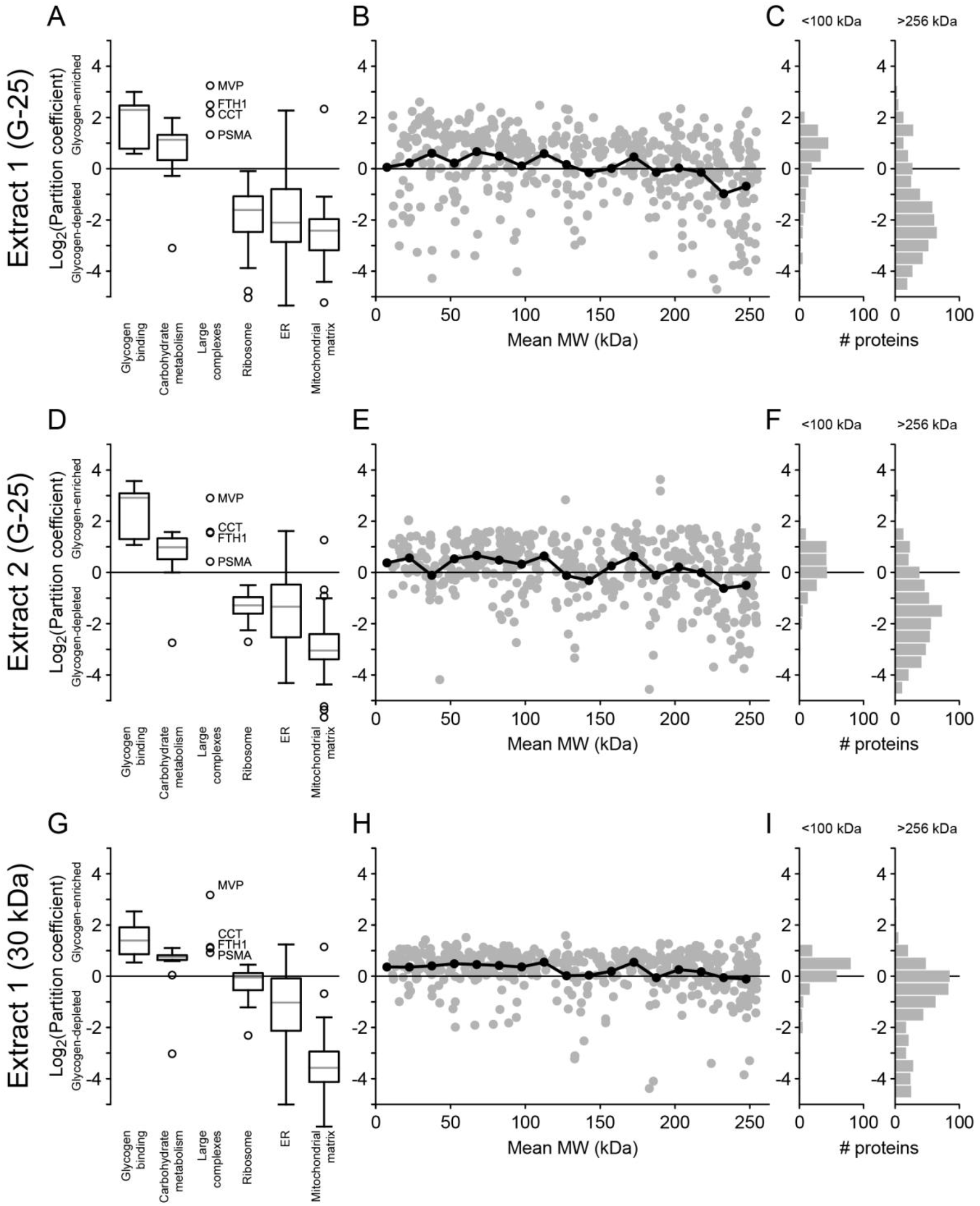
Replicate proteomics analyses suggest partitioning depends on binding and native MW. (Related to Fig 5) Extracts were crowded by two methods: (A-F) Sephadex G-25 gel filtration resin or (G-I) 30 kDa MWCO centrifugal filter unit. (A,D,G) Log base 2 partition coefficients of proteins sorted by Gene Ontology terms. Gray lines represent median values, boxes represent first and third quartile values, whiskers show the range of the data up to 1.5x the interquartile range, and circles represent outliers. (B,E,H) Log base 2 partition coefficients with respect to native MW less than 256 kDa. (C,F,I) Log base 2 partition coefficients of all proteins with native MW less than 100 kDa or greater than 256 kDa.

**Video S1. Liquid behavior of phases**. (Related to Fig 1D) 1.4x crowded extract was confined between coverslips then imaged immediately to observe spreading flow. Both phases exhibited liquid behavior, including deformation under flow, splitting, fusion, and rounding by surface tension. Extract was imaged by differential interference contrast (DIC) microscopy with a 20x objective.

**Table S1. Full proteomics data**. (Related to Figs 5, S2).

